# The unexpected provenance of components in eukaryotic nucleotide-excision-repair and kinetoplast DNA-dynamics from bacterial mobile elements

**DOI:** 10.1101/361121

**Authors:** Arunkumar Krishnan, A. Maxwell Burroughs, Lakshminarayan M. Iyer, L Aravind

## Abstract

**Background:** Protein ‘weaponry’ deployed in biological conflicts between selfish elements and their hosts are increasingly recognized as being re-purposed for diverse molecular adaptations in the evolution of several uniquely eukaryotic systems. The anti-restriction protein ArdC, transmitted along with the DNA during invasion, is one such factor deployed by plasmids and conjugative transposons against their bacterial hosts.

**Results:** Using sensitive computational methods we unify the N-terminal single-stranded DNA-binding domain of ArdC (ArdC-N) with the DNA-binding domains of the nucleotide excision repair (NER) XPC/Rad4 protein and *Trypanosoma* Tc-38 (p38) protein implicated in kinetoplast(k) DNA replication and dynamics. We show that the ArdC-N domain was independently acquired twice by eukaryotes from bacterial mobile elements. One gave rise to the ‘beta-hairpin domains’ of XPC/Rad4 and the other to the Tc-38-like proteins in the stem kinetoplastid. Eukaryotic ArdC-N domains underwent tandem duplications to form an extensive DNA-binding interface. In XPC/Rad4, the ArdC-N domain combined with the inactive transglutaminase domain of a peptide-N-glycanase originally derived from an active archaeal version, often incorporated in systems countering invasive DNA. We also show that parallel acquisitions from conjugative elements and bacteriophages gave rise to the Topoisomerase IA, DNA polymerases IB-Ds, and DNA ligases involved in kDNA dynamics.

**Conclusions:** We resolve two outstanding questions in eukaryote-biology: 1) origin of the unique DNA lesion-recognition component of NER; 2) origin of the unusual, plasmid-like features of kDNA. These represent a more general trend in the origin of distinctive components of systems involved in DNA dynamics and their links to the ubiquitin system.

## INTRODUCTION

Diverse selfish elements including bacteriophages, plasmids, and conjugative transposons possess the capacity for proliferation within the cell or the genome of their hosts. Thus, they are unceasingly entwined in multilevel conflicts with the host and other co-resident genetic elements, which possesses mechanisms to combat the negative effects of these entities to its own fitness [1-3]. Such inter-genomic and intra-genomic biological conflicts have spawned numerous molecular adaptations that function as “biochemical armaments” in both cellular genomes and the selfish elements: prime examples include restriction-modification [4, 5], toxin-antitoxin [6], CRISPR/Cas [7], and polyvalent protein systems [8] among others. Intriguingly, examination of some of these above-listed prokaryotic conflict systems has also led to the realization that they are potential evolutionary “nurseries” for molecular innovation spurred by the pressures for rapid adaptations. These adaptations are then disseminated via lateral transfer and used in functional contexts, which are very distinct from their original role in biological conflicts. Thus, we see numerous molecular adaptations unique to eukaryotes, such as enzymatic and DNA-binding domains in chromatin proteins [9-11], components of the RNAi systems and specialized components of the DNA-repair and -recombination having their ultimate evolutionary roots in prokaryotic conflict systems [1, 12-15].

The ArdC proteins from the plasmids pSA and RP4 are the founding members of a class of proteins, delivered by plasmids and certain phages into the host cell along with their DNA during invasion, termed “polyvalent proteins”, which at the interface of the biological conflict with their hosts [8, 16]. The ArdC protein has been shown to be an anti-restriction factor in the lncW plasmid pSA [16] and was also shown to bind single-stranded DNA (ssDNA) [16]. Of note, the ArdC protein is also known to be fused to the TraC-like primase domain as a component of the machinery involved in the conjugation-coupled replication of the plasmid RP4 [16-18]. Our recent study of counter-host strategies deployed by plasmids and phages showed that the classic ArdC protein of pSA contains two globular domains: a distinct N-terminal domain (ArdC-N) with the DNA-binding function and a C-terminal zincin-like metallopeptidase (MPTase) domain (Fig. 1A). The ArdC-N is one of the most prevalent domains in the polyvalent proteins and is coupled with multiple domains possessing an array of disparate effector activities or other DNA-binding domains [8] (Fig. 1A). Together, these observations suggested that ArdC might perform its anti-restriction function by coating ssDNA via the ArdC-N domain during invasion and also possibly by deploying the C-terminal MPTase domain to target the restriction endonucleases for cleavage or autoproteolytically releasing other effector domains coupled to the ArdC-N domain in polyvalent proteins [19].

**Fig. 1.**
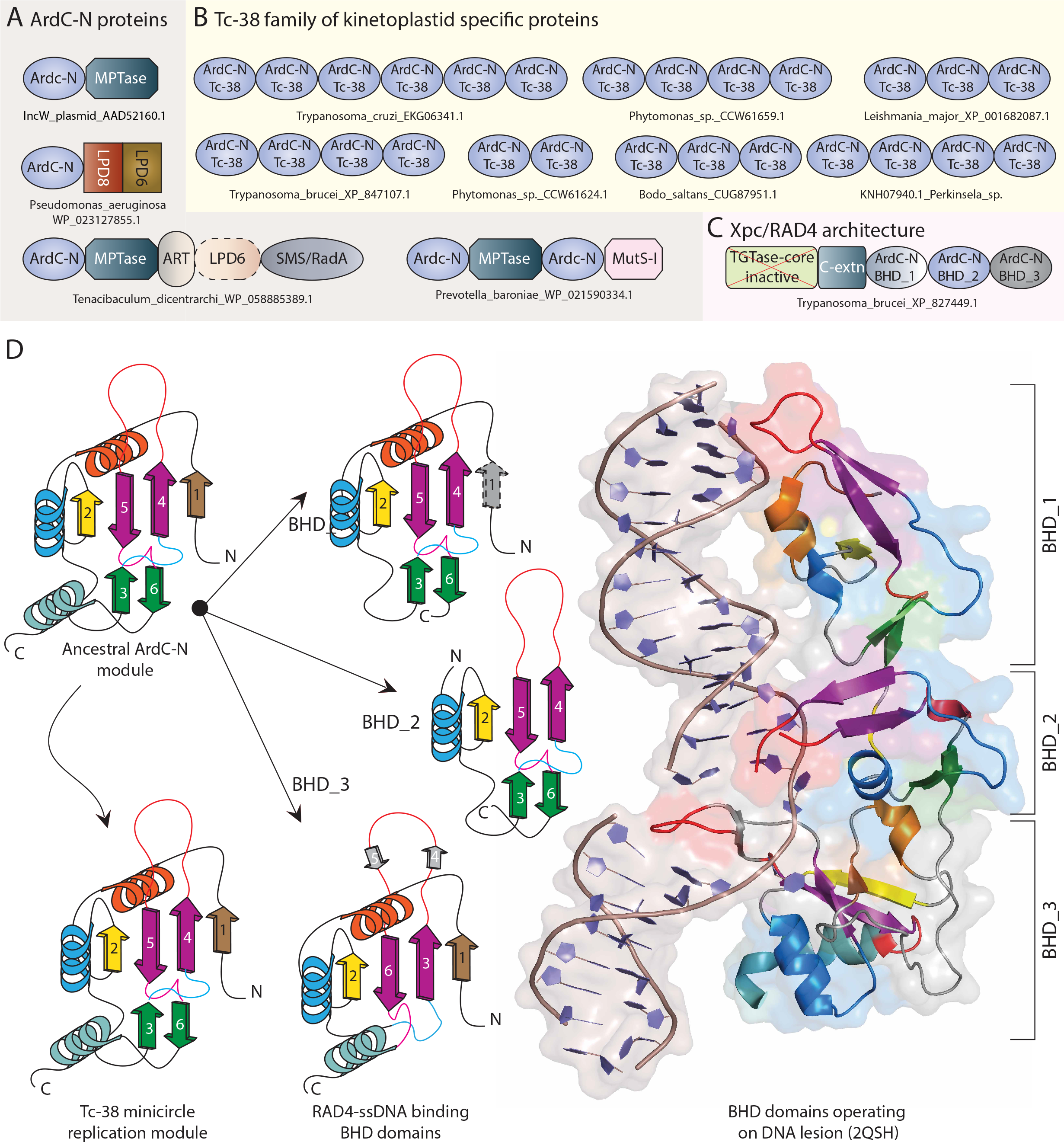
Domain architectures of a few exemplars of ArdC-N proteins (A), Tc-38 (B) and XPC/Rad4 (C) proteins. Proteins are labeled by their full species name and accession number. D) Schematic topological rendering on the left shows the ssDNA binding structural scaffold shared between the ancestral ArdC-N module, Tc-38 minicircle replication module and the XPC/Rad4 BHDs of the. The major β-sheet of this domain is formed by four β-strands (strands 1,2,4 and 5), while the β-strands 3 a nd 6 form the minor sheet: the characteristic long-hairpin loop connects the central two strands of the major sheet. Ribbon diagram on the right shows DNA-binding interface of t he BHD domains (PDBID: 2QSH): Secondary structure elements are colored the same as shown on the topological diagram (left). The long hairpin loops, is inserted into the DNA double helix in the recognition of the DNA-damage site in nucleotide excision repair.

In course of that study [8], we also detected significant sequence similarities between the ArdC-N domain and the *Trypanosoma* Tc-38 (p38) protein. In addition, our searches pointed to a potential evolutionary relationship between ArdC-N and the DNA-binding domains of the nucleotide excision repair (NER) XPC/Rad4 protein [20]. Tc-38 is a DNA-binding protein that associates with the structurally complex DNA network of the kinetoplastid mitochondrion known as kinetoplast DNA (kDNA) [21]. While kDNA exhibits considerable structural diversity across kinetoplastids leading to a classification scheme, which includes the so-called pro-kDNA, poly-kDNA, pan-kDNA, and mega-kDNA [22], in general, it consists of two recognizable classes of DNA “circles”: the maxi- and minicircles. Maxicircles are larger DNA rings (20 to 40 kbp) found in dozens of identical copies encoding rRNAs and cryptic genes coding for proteins involved in the mitochondrial energy transduction machinery [22-25]. In contrast, minicircles are a class of small DNA rings (0.5 to 2.5 kbp) displaying remarkable sequence heterogeneity found as several thousand copies per kDNA network and encode guide RNAs (gRNAs) that act as templates for directing RNA-editing via insertion or deletion of uridylate residues into the maxicircle-derived cryptic transcripts to yield functional messenger RNAs [22-25]. Tc-38 was shown to specifically function in ssDNA binding during replication and maintenance of kDNA, specifically influencing the count and supercoiling of minicircles [21, 26, 27].

Building on these observations, we detail herein the unification of the ArdC-N domain with the Tc-38-like and C-terminal DNA-binding domains of the XPC/Rad4 proteins. We trace the evolutionary trajectory of this newly-recognized DNA-binding fold and find that the ArdC-N module was horizontally transferred to eukaryotes from bacterial conjugative elements, likely twice independently. On one occasion, transfer of the ArdC-N played a role in the emergence of the lesion-recognition domains of XPC/Rad4 protein. On the other it was recruited for a role in kDNA-binding. This version of the ArdC-N domain underwent extensive expansion in the kinetoplastid lineage, possibly complementing the diversification/expansion of kDNA circles. We also show that the catalytically inactive transglutaminase domain of Rad4 emerged from ancestral archaeal PNGases and then fused with an ArdC-N from a bacterial plasmid giving rise to the extant form of XPC/Rad4. These findings throw new light on the provenance of certain eukaryotic systems that have thus far remained largely inscrutable. They also improve our understanding of the mechanism of eukaryotic nucleotide excision repair and kDNA replication in kinetoplastids.

## RESULTS AND DISCUSSION

### Identification and structural analysis of eukaryotic homologs of the ArdC-N domain

#### The Tc-38 family of kinetoplastid-specific DNA-binding proteins contains multiple copies of the ArdC-N domain

While investigating the ArdC-N domain in prokaryotes we initiated recursive sequence profile searches using the PSI-BLAST program against the non-redundant (NR) protein database of National Center for Biotechnology Information (NCBI). Initial iterations recovered the prototypical prokaryotic versions of these domains in the polyvalent proteins, which we discussed in detail in an earlier study [8]. These ArdC-N domains were almost always found at the N-termini of multidomain polypeptides of conjugative elements across bacterial lineages and sporadically in euryarchaeota and nanoarchaeota [8]. However, we surprisingly also recovered proteins with this domain from eukaryotes albeit with domain architectures completely unlike those of the prokaryotes. For example, a search initiated with an ArdC-N domain from *Salmonella enterica* against the NCBI NR database (WP_023226849.1: residues 1 to 140) recovered a significant relationship with Tc-38-like ssDNA-binding protein from the deep-branching kinetoplastid *Perkinsela* sp. (KNH05906.1 (e-value: 3e-06); KNH07778.1 (3e-04); KNH08381.1 (3e-04) in iteration 2 with max target sequences set to 20000) and those from crown-group kinetoplastids such as *Trypanosoma* vivax (CCC49616.1). A reciprocal search using Tc-38-like from *Perkinsela* (KNH07778.1) easily recovered the TC-38 family proteins from other kinetoplastids and several bacterial ArdC-N homologs with e-values reaching 1e-05 in PSI-BLAST iteration 2. This affirmed the presence of an ArdC-N domain in Tc-38. Subsequently, multiple searches using diverse Tc-38 sequences as search seeds successively recovered further related sequences from the kinetoplastids, pointing to a large expansion of Tc-38-like ssDNA-binding proteins from diverse kinetoplastids including early-branching representatives such as *Perkinsela* sp and *Bodo saltans* (see exemplars in Fig. 1B). Tailored searches for further versions in other euglenozoans such as the diplonemids and euglenids failed to recover reliable homologs of Tc-38, thus suggesting that Tc-38 are specific to kinetoplastids (Supplementary Table S1 at ftp://ftp.ncbi.nih.gov/pub/aravind/ardcn/ardcn.html)

#### The so-called BHD domains of nucleotide excision repair (NER) protein Rad4/XPC are ArdC-N domains

To further investigate the ArdC-N domain, a Hidden Markov Model (HMM) profile constructed from a multiple alignment of ArdC-N sequences was searched against a database of HMM profiles constructed from the Pfam database [28] and individual Protein Databank (PDB) [29] entries (see Methods) with the HHpred program [30]. These searches surprisingly detected a significant relationship between the ArdC-N domain and the Pfam profile “BHD_2”, one of three domains labeled BHD hitherto exclusively observed in the C-terminal DNA-binding region of the nucleotide excision repair proteins XPC/Rad4 (p-value: 2.2E-08, probability: 96.7%; PDB-ID: 2QSH [20], p-value: 8E-07, probability: 93.8%). Reverse profile-profile searches of the sequences corresponding to the “BHD_2” model from Pfam recovered not just the eukaryotic XPC/Rad4 proteins but also bacterial exemplars of the ArdC-N domain. For instance, an iterative sequence profile-based search initiated with a BHD_2 sequence from the fungus *Candida albicans* against the NCBI NR database (XP_712990.1; residues 446 to 507) retrieved the ArdC-N domain in *Fictibacillus phosphorivorans* (WP_066238349.1, e-value: 1e-06, iteration: 5), *Bacillus oceanisediminis* (WP_019379886.1, e-value: 2e-04, iteration: 5), *Bacillus* sp. (WP_009336457.1, e-value: 3e-04, iteration: 5), among others. The BHD_2 is the central domain in the tripartite organization of the C-terminal DNA binding region of XPC/Rad4, flanked N-terminally by the BHD_1 and C-terminally by the BHD_3 domains [20]. N-terminal to the BHD_1-3 module, the XPC/Rad4 proteins are further fused to an inactive transglutaminase fold domain (see following sections) [31] (Fig. 1C).

Visual inspection of the individual BHD domain structures and analysis of concordance in secondary structure elements strongly suggested a relationship across the three domains (Fig. 1D). Accordingly, pairwise structural homology searches (see Methods) initiated with the discrete domains as queries were confirmatory, consistent with previous studies suggesting a possible relationship between the three BHDs [8]. For example, a pairwise search initiated with the BHD_1 domain recovers the region corresponding to the BHD_2 domain with scores strongly indicative of a relationship (Z-score: 4.6). The reverse search initiated with BHD_2 likewise recovers BHD_2 (Z-score: 7.2) and BHD_1 (z–score: 3.6). Thus, the above observations expand the range of established ArdC-N domains beyond the classical plasmid-interacting prokaryotic ArdC-N family to include the Tc-38-like and BHD families from eukaryotes.

#### Shared structural core and conserved features of the ArdC-N domain

Based on the multiple sequence alignments constructed using representatives of the ArdC-N domain from diverse taxa, we investigated the structural scaffold of the ArdC-N domain, specifically informed by the crystal structures of the versions found in the Rad4 protein (Fig. 1D and Fig. 2; also see Supplementary alignments S1-S4). These structure-informed alignments together with structure predictions indicated that the ancestral core of the ArdC-N domain is a rather distinctive structure with no close relationship to any other protein fold. These observations also dispel a previously held view that the BHDs in XPC/RAD4 were OB-fold domains [32, 33]. The ArdC-N domain is characterized by a couple of β-sheets: the major one formed by up to 4 strands and the minor one by two strands. The polypeptide chain crosses over at the point of entry and exit to the central two strands of the major sheet and the cross-over is bounded by the two strands of the minor sheet (Fig. 1D). Further, the cross-over region has a distinctive meander at the N-terminus and a single helical turn at the C-terminus which we term the squiggle (Fig. 1D). Together, these sheets form an open barrel-like structure. Despite the striking structural conservation, distinct ArdC-N clades display high sequence heterogeneity (Fig. 2; Supplementary alignments S1-S3). Nonetheless, one subtle yet notable conserved sequence signature across the ArdC-N domains marks the aforementioned squiggle: a hhsxxQ motif (with ‘h’ representing a hydrophobic residue, ‘s’ representing a small residue, and ‘x’ representing any residue; the first hydrophobic residue is usually aliphatic, while the second is aromatic) (Fig. 2A and B; Supplementary alignments S1-S2). The glutamine that marks the end of the motif and occurs just prior to the second strand of the second sheet is mostly conserved; notable exceptions being BHD_1 and BHD_3 (Fig. 2C and D; Supplementary alignments S1-S3). This glutamine appears to play a role in maintaining the distinctive structure by stabilizing the squiggle via a contact with the backbone. Further, comparable squiggles observed in other, distinct protein folds have been linked to regions displaying conformational flexibility in a fold [34-36]. The ArdC-N squiggle and the accompanying conserved glutamine residue could similarly facilitate conformational change during recognition of specific features in DNA.

Diverse ArdC-N domains often retain certain aromatic residues previously implicated in mediating DNA contacts [32]. A prime example is a conserved hydrophobic position (most frequently a tryptophan) observed at the beginning of helix-2 (helix-1 for BHD_2) across most versions of the domain (Fig. 2). The BHD versions of the ArdC-N domain in XPC/RAD4 further indicate that different representatives of the domain might potentially adopt distinct modes of binding DNA[20]. However, a common denominator is the contact made with single-stranded or double-stranded DNA by the long-hairpin loop connecting the central two strands of the major sheet (Fig. 1D). Structural studies on the Rad4 protein implicate the deep insertion of this loop into the DNA double helix in the recognition of the DNA-damage site in NER [20] (Fig. 1D). Further, version-specific contacts with the DNA-backbone are mediated by the helices downstream of the first and second strands of the core (Fig 1D).

**Fig. 2.**
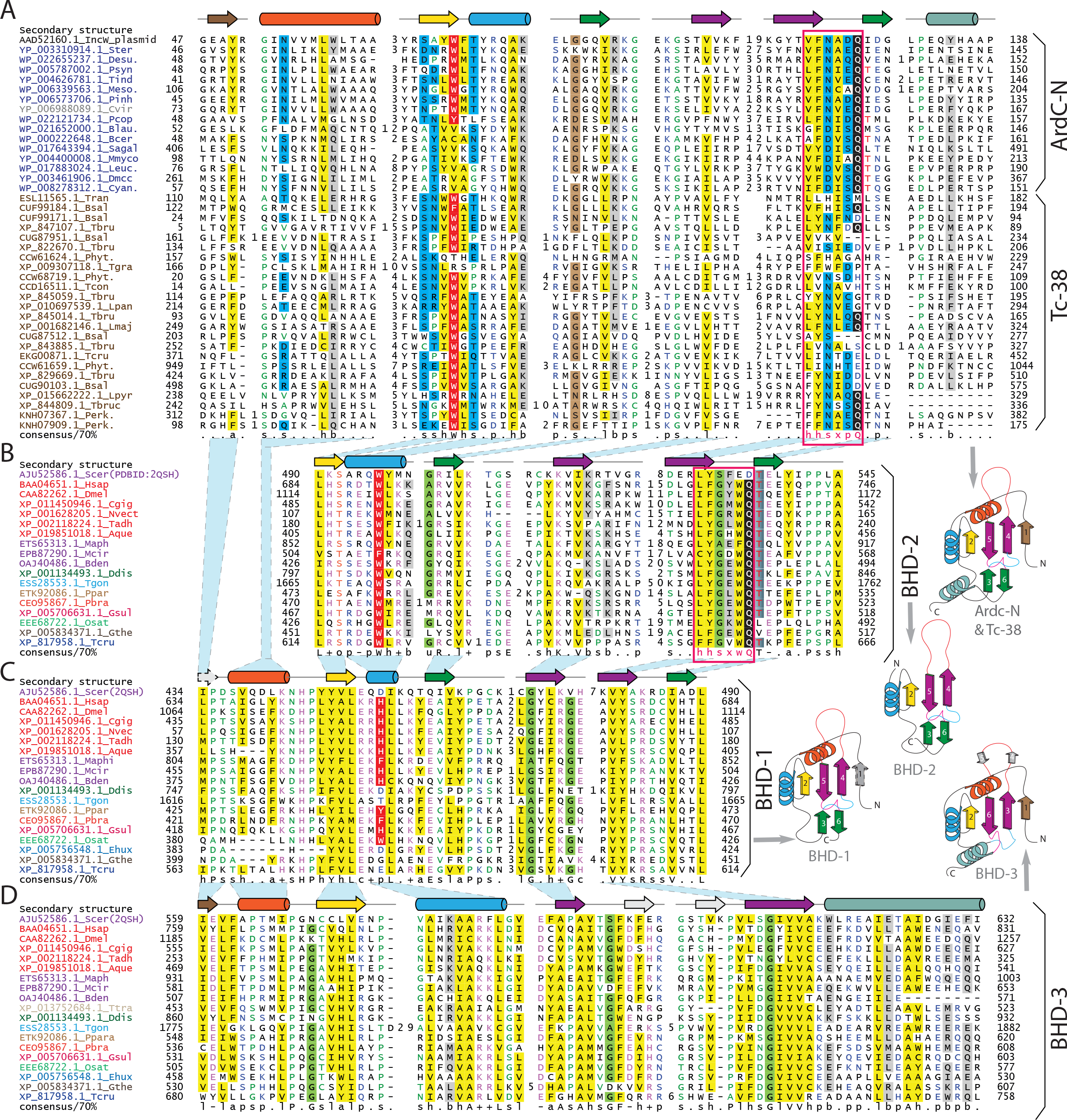
Multiple sequence alignments of ArdC-N/Tc-38 (A), BHD2 (B), BHD1 (C) and BHD3 (D) domains. Secondary structure elements are shown on the top and colored the same as show n in topological diagrams shown to the right. The characteristic hhsxxQ motif is highlighted as red rectangular box. Polar and small residues shared between the ArdC-N and Tc-38 to the exclusion of other domains are highlighted in blue and light brown, respectively. Conserved aromatic residues (typically a tryptophan, phenylalanine and histidine) are highlighted in red.

### Unraveling the complex evolutionary history and functional implications of proteins containing the ArdC-N domain in eukaryotes

#### Eukaryotes acquired the ArdC-N domain through two independent transfers from bacteria

Notably, searches initiated with Tc-38-like or BHD family members never directly recover each other as immediate hits but recover different bacterial ArdC-N domains as their best hits. This observation is further consistent with: (i) distinct phyletic distribution of BHD domains and Tc-38: Tc-38 proteins are restricted to the kinetoplastids (Supplementary Table S1), whereas the BHD-domain-containing XPC/Rad4 family is widespread across eukaryotes (Supplementary Table S2); (ii) subtle yet clearly distinct conservation patterns shared by the bacterial ArdC-N respectively with the BHDs and Tc-38 families to the exclusion of the other: the bacterial ArdC-N and BHD_2 specifically share a threonine residue immediately following the conserved Q of the hhsxxQ motif, while in Tc-38 the equivalent residue is mostly hydrophobic (Fig 2A and B; Supplementary alignments S1 to S3). Conversely, the ArdC- N domains in Tc-38 domains share several features establishing a close relationship with the bacterial ArdC-N domains to the exclusion of the BHDs: (i) clear conservation of both N-terminal α-helices containing shared polar residues (typically an asparagine, serine, and threonine), located at the beginning of helices 1 and 2 and strand-2 (Fig. 2A; Supplementary alignments S1-S3); (ii) similarly, shared polar residues, typically a serine, aspartate, or glutamate, as well as an asparagine, were found upstream of the characteristic Q residue (Fig. 2A) (iii) strong conservation of a small residue, typically a glycine, between the second conserved helix and the β-strand. These observations suggest that the prokaryotic ArdC-N domains were acquired independently on two distinct occasions by the eukaryotes: one of these acquisitions led to the more broadly distributed BHD versions found in the XP-C/Rad4 proteins, whereas the second acquisition led to the kinetoplastid-specific Tc-38-like versions. To better understand this dual acquisition of the eukaryotic ArdC-N domains we next systematically investigated the evolutionary histories of the XP-C/Rad4 and TC-38-like proteins and explored the potential functional implications of their constituent domains.

#### The origin of XPC/Rad4 through the confluence of domains with distinct evolutionary histories acquired from archaea and conjugative elements

More than a decade ago, we had shown that XP-C/Rad4 contains an inactive copy of the transglutaminase (TGL) domain, an ancient version of the papain-like peptidase fold [31]. This domain occurs N-terminal to the ArdC-N domains (BHD_1:3) described above. Further, this TGL domain is specifically related to the catalytically active version found in the Peptide-N-glycanase (PNGase) and the two are unified by a conserved C-terminal extension to the exclusion of other members of the TGL superfamily [31]. To elucidate the origin of this architectural linkage between the TGL and ArdC-N domains in XPC/Rad4 we surveyed the new genomic data available since our earlier study relating to the TGL domain. Iterative PSI-BLAST searches recovered sequences from thaumarchaeota (ALI37408.1; OLE40692.1; OLC36996.1) as the closest-related non-eukaryotic homologs of the XP-C/RAD4-PNGase TGL domain. Using these sequences as search seeds, we recovered additional homologs of archaeal PNGases from Euryarchaeota and Candidatus Micrarchaeota belonging to the DPANN group of archaea; further searches with these recovered homologs from other archaeal lineages including the Crenarchaeota (Supplementary Table S3). We then used these sequences together with the eukaryotic XP-C/Rad4, and PNGases to construct a phylogenetic tree based on their shared TGL domain. Earlier identified related TGL domains found in ky/cyk3 and YebA [31] served as outgroups in this analysis. Saliently, the tree showed that that the TGL domains from Thaumarchaeota are the closest sister group of the eukaryotic XP-C/Rad4 and PNGase TGL domains (Supplementary Data S1: tree files). In support of this specific relationship, the thaumarchaeal versions share with eukaryotic PNGases a unique Zn-binding domain N-terminal to the TGL domain (Fig. 3A). In contrast to the thaumarchaeal and eukaryotic versions, all other archaeal PNGase homologs are predicted cell-surface proteins as indicated by their N-terminal signal peptide and C-terminal transmembrane helix (Fig. 3A). This is consistent with the recent proposal that the archaeal progenitor of the eukaryotes was derived from within an assembly of archaea which includes the Thaumarchaeota and the Asgardarchaeota [37].

**Fig. 3.**
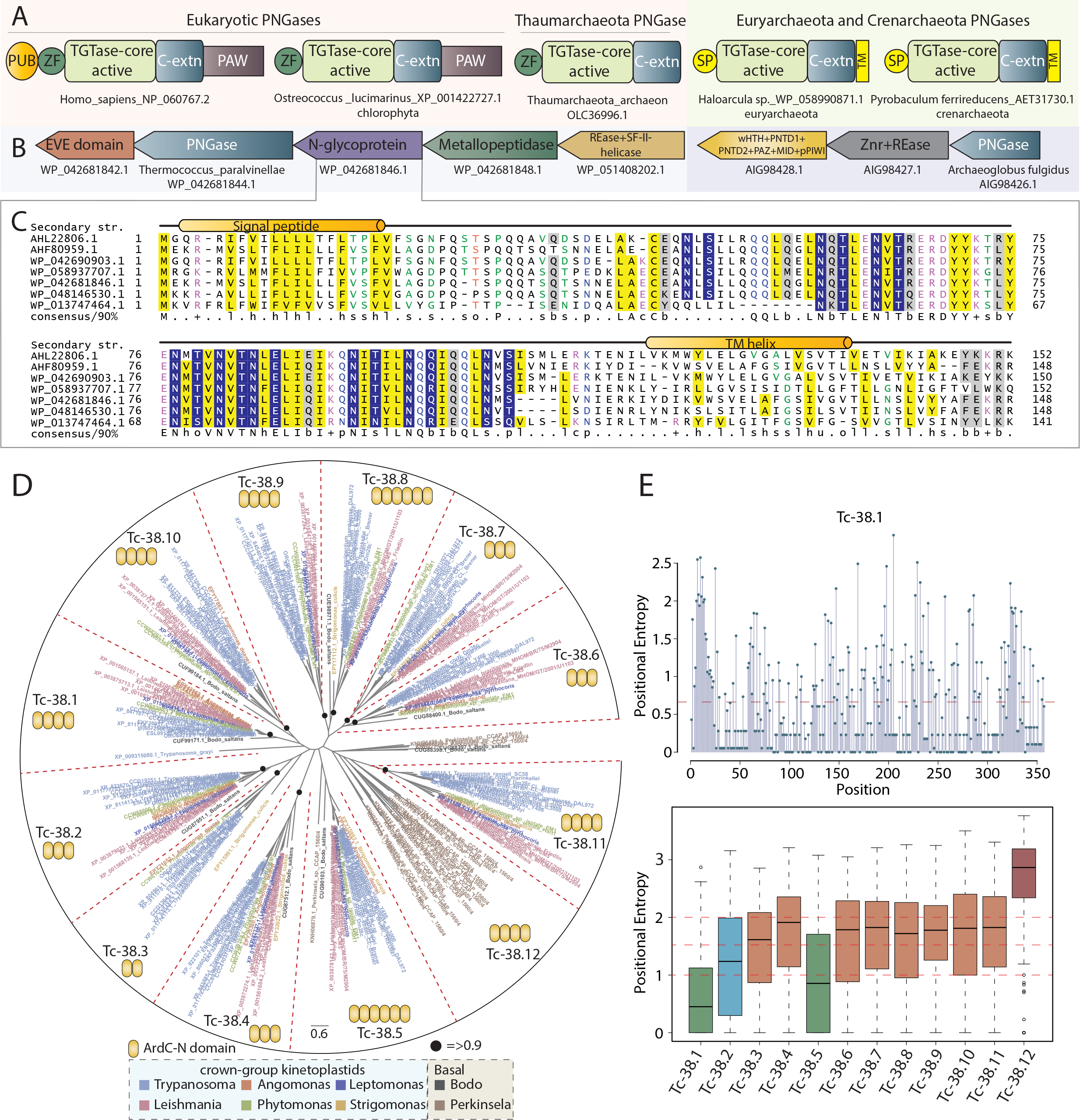
A) Domain architectures of archaeal and eukaryotic PNGases. B) Gene neighborhoods of PNGases from *Thermococcus paralvinellae* (WP_042681844.1) and *Archaeoglobus fulgidus* (AIG98426.1) are shown. The N-glycoprotein, a substrate for the active PNGase for de-N-glycosylation is operonically linked. C) Multiple sequence alignment of the predicted N-glycoprotein showing characteristic NxT/S and NxE/Q repeats (highlighted in deep blue). D) Phylogenetic tree showing th e Tc-38 family expansions grouped into 12 distinct clusters. *Perkinsela*. *sp* encoding a lineage-specific expansion is highlighted in brown. The number of ArdC-N domains present within the same polypeptide for each distinct cluster are shown as insets (domains colored yellow). E) Positional entropy values for the slow-evolving Tc-38.1 cluster with mean entropy value (dotted horizontal line in red). Further a box plo t is shown comparing global entropy values for the 12 distinct Tc-38 clusters. Mean entropy values of >2, >1.5, and <1.5 and <1 are colored deep-brick, light-brown, light-blue, and green, respectively.

Analysis of the gene neighborhoods (Fig. 3B; Supplementary Table S3) of the archaeal versions revealed that in several euryarchaea the PNGase homolog is coupled with a gene coding for a predicted cell-surface protein of 160-180 amino acids with 6 conserved repeats containing an NxT/S motif and two additional NxQ/E motifs (Fig. 3C). Such repeats with conserved asparagines suggest that this novel polypeptide is an N-glycoprotein and is the likely substrate for de-N-glycosylation by the associated PNGase homolog. Sequence alignments demonstrated that like the eukaryotic PNGases, but unlike XPC/RAD4, all archaeal homologs, including thaumarchaeota, retain an intact catalytic triad suggesting that they are active enzymes (Supplementary alignment S5), which have the capacity to catalyze a PNGase-like reaction. Together these observations indicate that the archaeal PNGase homologs are likely to function similar to the eukaryotic PNGases. Interestingly, these archaeal PNGases also show further linkages to genes encoding proteins with an EVE domain with the PUA-like fold implicated in the recognition of modified bases in DNA [38], a REase (restriction endonuclease) fold DNase domain and in some cases a PIWI family protein involved in RNA-dependent restriction [12, 39-41] (Fig. 3B). Hence, these gene-neighborhoods are likely to code for systems potentially involved in the discrimination and restriction of invasive elements with DNA containing modified bases, similar to other such biological conflict systems with comparable components that are widespread across prokaryotes [19, 38, 42]. However, the noteworthy twist in these systems is the inclusion of a cell-surface PNGase and/or a cell-surface N-glycosylated glycoprotein component. Therefore, these appear to be dual action systems featuring cell-surface components interacting with the invasive elements in the extracellular compartment to potentially preclude their entry via de-glycosylation of surface receptors and associated restriction-components that would target their DNA intracellularly.

PNGases are present across all sampled major eukaryotic lineages and XPC/Rad4 is present in most except for the diplomonads, breviatea, and centroheliozoa (Supplementary Table S4). Among the basal eukaryotic lineages, parabasalids (e.g. *Trichomonas*) possess a second paralogous PNGase-like protein, which, like XPC/Rad4, is inactive but lacks an association with the ArdC-N domains. The earliest occurrences of the inactive TGL together with the ArdC-N domains are found in euglenozoans and heteroloboseans (Supplementary Table S4). These observations imply that the ancestral eukaryote likely possessed a PNGase-like protein inherited from the archaeal progenitor. Early in eukaryotic evolution, this appears to have spawned two paralogs, one retaining the catalytic activity similar to the ancestral PNGase and the second becoming inactive as seen in the parabasalids. After the basal-most eukaryotic lineages such as parabasalids and diplomonads branched off, this inactive version appears to have fused with the ArdC-N domain acquired from a bacterial conjugative element resulting in the core architecture of the extant XPC/Rad4. Within the context of the XPC/Rad4, the ArdC-N domain underwent a triplication. Of the three copies, the BHD_2 version remained closest to the plasmid ArdC-N at the sequence level but lost certain structural elements (Fig. 1D and Fig. 2B). Conversely, the BHD_1 and BHD_3 versions have diverged considerably at the sequence level while retaining most core structural features (Fig. 1D and Fig. 2C and D).

#### The evolutionary history of the XPC/Rad4 explains the unique aspects of eukaryotic NER

The above-reconstructed scenario allows us to explain the conundrum of the unique components of the NER system found in eukaryotes. NER, which is required for removal of bulky and helix-distorting alterations of DNA [43] (e.g. thymine dimers) is found across the bacterial and archaeo-eukaryotic branches of life. In both the branches of life, the mechanistic details are comparable with distinct components that respectively recognize the DNA lesion, catalyze local unwinding of the DNA duplex and the incision endoDNases, which cut out the ssDNA segment with the lesion [44]. However, these components are unrelated in the bacterial and the archaeo-eukaryotic branches of life. In the former, it is mediated by the UvrABC complexes [45], whereas in the latter XPB/Rad25 and XPD/Rad3 helicases catalyze the unwinding activity and XPF/ERCC1 functions as one of the conserved incision endoDNases [46-49]. However, the lesion-recognition component is not conserved between archaea and eukaryotes. Studies in archaea have implicated the SSB (RPA) protein in this role, which also has an important function in binding ssDNA during replication initiation [50]. Hence, our finding that early in eukaryotic evolution, the ArdC-N domains were recruited for the lesion recognition role indicates the that an alternative protein with comparable ssDNA recognition capacity to SSB(RPA) displaced it. Notably, our analysis also shows that it was likely acquired by the eukaryotes from the replication system of a plasmid/conjugative transposon probably borne by a bacterial endosymbiont. This might suggest that the ssDNA replication intermediates of these conjugative elements were better structural analogs of the DNA lesions that trigger NER in eukaryotes and the ArdC-N domain was accordingly better equipped to recognize them than the ancestral SSB(RPA).

In eukaryotic NER other proteins hr23B/Rad23B, centrin-2, DDB1, DDB2, and cullin-4 augment DNA-lesion-recognition by XPC/Rad4 either by increasing its stability, binding efficiency and/or increasing the range of distinct DNA adducts that are recognized [51-55]. Some of these, namely DDB1, DDB2, and cullin-4, emerged relatively late in eukaryotic evolution via duplication of more widely-distributed paralogs (AM Burroughs, L Aravind, personal observations). Centrin-2 and hr23B/Rad23B are present across eukaryotes including the basal lineages and are involved in more general functions [56-61]. This points to a recruitment to NER from these more general, ancestral functions. Most notable among these is the hr23B/Rad23 which contains a ubiquitin-like domain, an XPC/Rad4-binding (R4BD) domain, and two ubiquitin-associated (UBA) domains. Interestingly, the R4BD interacts with the TGL domain of both XPC/Rad4 and PNGase [62, 63]. In this context, our findings also help explain how the other module of XPC/Rad4, the TGL domain, was recruited for the NER from a protein originally involved in glycoprotein degradation (PNGase). The shared architectural features of thaumarchaeal and eukaryotic PNGases suggests that an enzyme for the intracellular degradation of misfolded N-glycosylated glycoproteins likely emerged in their common ancestor from an earlier archaeal PNGase-like cell-surface protein involved in deglycosylation of cell-surface glycans. By the time of the ancestral eukaryote this was coupled with ubiquitin-mediated protein degradation through the 26S proteasomal system, to which the PNGase is recruited by a complex formed with hr23B/Rad23 [62-64]. The similar Rad4·Rad23 complex formed via the inactive TGL domain instead protects XPC/Rad4 from proteasomal degradation and allows it to more efficiently complete NER [61]. Given that the ArdC-N domain of the bacterial conjugative elements is also most frequently fused to an active or inactive peptidase domain (Fig. 1A) [8], it is probable that version first acquired by the eukaryotes not just recognized DNA lesions but also mediated other interactions via this peptidase domain. This was likely displaced by the inactive peptidase-like TGL domain from the PNGase paralog, which now allowed a comparable interaction with hr23B/Rad23 and its sequestration for stabilizing XPC/Rad4 against proteasomal destruction. This opposing regulation via proteasomal degradation and hr23B/Rad23 stabilization was probably selected because it allowed a threshold-based regulation of the DNA-incision activity of NER, which if unregulated might result in unnecessary and deleterious DNA breaks.

#### The recruitment of ArdC-N in kinetoplastids and its subsequent lineage-specific expansions

As noted above, our iterative searches recovered a previously-unrecognized expansion of Tc-38-like proteins with ArdC-N domains in kinetoplastids. We failed to identify any Tc-38-like proteins from other major classes of euglenozoans such as the Diplonemida, Euglenida, and Symbiontida. However, it should be noted that the sequence data from the other euglenozoans is currently limited. Two distinct classification methods indicated that these Tc-38-like proteins form 12 clades of paralogs (see Methods). We name these clades Tc-38.1 to Tc-38.12 (see Supplementary alignments S4). Eleven of these (Tc-38.1-11) contain members from the crown kinetoplastid lineage, the Trypanosomatidae (*Trypanosoma*, *Leishmania*, *Leptomonas*, *Phytomonas*, *Angomonas*, *Strigomonas*) (Fig. 3D). Further, seven of these 11 paralog clades could be traced back to the more basal *Bodo saltans*, with the versions from this organism emerging as the basal-most member of each of those clades in the phylogenetic tree (Fig. 3D). Two additional Tc-38-like proteins in *Bodo* did not unambiguously associate with any of these paralogous clades. The 12th clade of TC-38-like proteins is exclusively comprised of a large LSE of 30 members from the basal kinetoplastid *Perkinsela* (Fig. 3D). Hence, the most plausible evolutionary scenario would be that the representatives of at least seven of these paralogous clusters were already present in the last common ancestor of Bodonidae and Trypanosomatidae, suggesting an early diversification via LSE resulting in the founding members of the paralogous groups, somewhere close to the divergence of Bodonidae from *Perkinsela*. Given that the other euglenozoans apparently lack homologs of Tc-38, the second currently known acquisition of ArdC-N domains in eukaryotes likely happened at the base of kinetoplastida through the transfer of an ArdC-N domain from a plasmid/conjugative element (see below) probably residing in an endosymbiotic bacterium. In course of the expansion of this protein family during kinetoplastid evolution we observe 1) repeated duplications of the ArdC-N domain and in some cases losses of the ArdC-N domain within the same polypeptide. Thus, these paralog clades are characterized by anywhere between two to six copies of ArdC-N domains, with three or four repeats being most frequent (Fig 3D and Supplementary alignments S4). 2) Extreme sequence diversification of the individual ArdC-N domain repeats both within the same polypeptide and across members from the same paralog clade. 3) Structural divergence within the loop-region in the central β-hairpin of the domain and in some cases expansions of long regions of low-complexity linking adjacent ArdC-N domains (Supplementary alignments S4).

#### Evidence for Tc-38-like proteins playing a key role in kDNA replication in kinetoplastids

Our analysis of the diversification of the Tc-38 family also throws light on their functions in the kDNA replication system and the diversification of kDNA structures. The universal minicircle sequence (UMS) sequence with a G-quartet at the start (e.g. UMS in *T. brucei* reads GGGGTTGGTGTA), is characteristic of the minicircle replication origins [22, 65, 66]. Tc-38 was shown to have a specific preference for binding TG-rich repeat regions comprising the UMS [27]. Moreover, it also binds the hexamer, a sequence found on the lagging template strand at the start of the first Okazaki fragment. However, a similar UMS-binding function has also been attributed to another protein named the universal minicircle binding protein (UMSBP) [67, 68]. This has led to the debate as to whether Tc-38 or UMSBP is the primary ssDNA binding protein in the kDNA replication system. We present several lines of evidence based on our analysis of the Tc-38-like proteins and the available experimental data that have a bearing on which of these is the principal minicircle origin recognition protein. First, Tc-38 like proteins are specific to kinetoplastids, as would be expected for a protein with a kDNA-specific function. However, UMSBP belongs to a family of proteins with the Zn-knuckle that are distributed throughout eukaryotes and implicated in functions such as ssRNA-binding in various RNA-processing systems like the mRNA maturation and splicing apparatuses [69]. Specifically, UMSBP does not exhibit the diversity and the expansions of the Tc-38-like family, complementing the diversification of kDNA circles in kinetoplastids (see below). Second, while the ArdC-N domain binds ssDNA in the related context of bacterial conjugative element replication intermediates, the CCHC-type zinc (Zn)-knuckle repeats found in UMSBP are known to be promiscuous in binding single-stranded nucleic acids and G-quartet structures [70]. Indeed, UMSBP, in addition to binding the template strand of the UMS, also quite promiscuously binds the complementary strand to the hexamer instead of the template strand [71]. Third, although the RNAi knockdowns of UMSBP led to defects in kDNA circle replication the primary effects are seen in segregation of daughter networks with additional effects on nuclear division [72]. Tc-38 is enriched in the mitochondrial fraction across most stages of the kinetoplastid life cycle and its RNAi-knockdown results in loss of kinetoplast DNA and accumulation of “free” minicircles detached from the central network known as the fraction S minicircles [21, 27]. Together, these observations favor Tc-38 being the primary UMS-associated circle replication origin-binding protein, with UMSBP likely playing a more general role in ssDNA binding in the kinetoplast and elsewhere.

In functional terms, based on the model of the triple ArdC-N domains in XPC/Rad4 [20] we propose that the Tc-38-like proteins in kinetoplastids can bind a considerable DNA stretch encompassing up to 24 base pairs [20] or more associated with the minicircle replication-origins. Moreover, in parallel to the ArdC-N domains in prokaryotic conjugative elements, Tc-38 likely acts as a ssDNA-protecting agent[26]. Tc-38-like proteins probably protect the integrity of the kDNA circles by providing a bulwark against supercoiling and enabling polymerase procession [21] similar to the role of ArdC-N during the transfer of the ssDNA replication intermediate of a conjugative element to a new host. These observations suggest that Tc-38 binds single-stranded minicircle DNA following initial helicase and topoisomerase activity, stabilizing the unwound structure during the initiation of leading and lagging strand synthesis. Loss of Tc-38, therefore, results in a continuation of the unwinding without initiation of synthesis [21]. Other observations point to a more pervasive role for the diversified repertoire of Tc-38-like proteins in kinetoplastids paralleling the evolutionary diversification of kDNA. We observed varying degrees of sequence divergence across the Tc-38 paralog clades ranging from well-conserved slow-evolving clades to rapidly diversifying clades (Fig. 3E). For instance, Tc-38.1, which contains the minicircle replication origin binding Tc-38 protein (also known as p38), is strongly conserved across both bodonids and trypanosomatids (as indicated by a low entropy value) (Fig. 3E). Such conservation indicates a selective pressure to likely both preserve the binding features of the Tc-38.1 and the corresponding sequence determinants recognized by it. However, other paralogous clusters display considerable sequence diversity (Fig. 3E). Notably, the knockdown of the Tc-38 gene by itself does not appear to affect all DNA circles [21] suggesting that some of the paralog clades have probably been optimized for recognition of sequences in specific groups of circles. Further, a great diversity of distinctive kDNA network structures that have been described across kinetoplastids. For instance, kDNA network structure observed in different basal kinetoplastids include: Pro- and Poly-kDNA, where minicircles exist as monomeric units and often are covalently closed and topologically relaxed; Pan-kDNa where minicircles are mainly monomeric, but unlike Pro- and Poly-kDNA, they form a supercoiled structure; Mega-kDNA, an unusual form of kDNA where they do not constitute actual minicircles, but minicircle-like sequences are tandemly linked to larger maxicircle-like molecules [22]. Thus, the observed ArdC-N domain diversity in the Tc-38-like family might have gone hand-in-hand with the emergence of diverse kDNA network structures.

#### Tc-38 is joined by several other components of the kDNA replication apparatus in being derived from selfish replicons

To contextualize our observation concerning the provenance of the Tc-38-like proteins we expanded our analysis of other kDNA replication components. Importantly, we observed that in a phylogenetic tree the kinetoplastid topoisomerase IA is lodged within a clade, which is otherwise mostly comprised of topoisomerases from bacterial plasmids/conjugative elements that also code for an ArdC-N domain protein (e.g. *Helicobacter cetorum* plasmid topo IA, Genbank: AFI04135.1; Fig. 4A). Strikingly, these mobile-element-topoisomerases are linked in a conserved gene-neighborhood with the gene for the ArdC-N domain protein as their immediate downstream neighbor (e.g. *Helicobacter cetorum* plasmid ArdC-N, Genbank AFI04134.1) (Fig. 4A). This suggests that both the topo IA and the ArdC-N precursor of the Tc-38 protein were likely to have been acquired from a common plasmid source and recruited together for kDNA replication. The primary kinetoplast DNA polymerases are members of the bacterial pol I family. In the phylogenetic tree, the four paralogous kinetoplastid enzymes Pol IB, IC, and ID formed respective clusters with internal branching largely in agreement with kinetoplastid phylogeny; the homologs from bacteriophages-infecting gammaproteobacteria are placed basal to these (Fig. 4B). This indicates the acquisition of the DNA pol I from a bacteriophage likely occurred in the common ancestor of the kinetoplastids and subsequently underwent multiple rounds of duplication giving rise to the four paralogs found across all kinetoplastids. Trypanosomatids possess two ATP-dependent DNA ligases k-α and k-β that are involved in kDNA replication. Interestingly, our sequence profile searches recovered a third divergent and previously unreported kinetoplastid-specific ATP-dependent ligase in both trypanosomatids and *Bodo saltans* (e.g. Genbank: CUI15152.1), which in searches recover ligases k-α and k-β, suggesting a specific relationship to the known kinetoplast DNA ligases (Fig. 4C). While these could be traced back to *Bodo*, the homologs from *Perkinsela* are placed basal to the overall clade separating k-α and k-β, suggesting the emergence of the three distinct ligase clades happened in the common ancestor of trypanosomatids and *Bodo* (Fig. 4C). In turn, all kinetoplast DNA ligases form a clade with those from gammaproteobacterial bacteriophages to the exclusion of other DNA ligases (Fig. 4C). These results emphatically established that at least two groups of key kDNA replicative enzymes, the DNA pol I and the ATP-dependent DNA ligases, were acquired from gammaproteobacterial bacteriophages. Aside from these, a member of the archaeo-eukaryotic primase superfamily PPL1, which functions as both a primase and a polymerase (primpol) has been proposed to be involved in both mitochondrial and nuclear DNA replication [73]. We have earlier demonstrated all eukaryotic primpols were ultimately inherited from a nucleo-cytoplasmic large DNA virus (NCLDV)-like source [13, 74]. Thus, several key kDNA replication components appear to have been acquired from not just plasmids/conjugative elements but also bacteriophages and eukaryotic viruses.

**Fig. 4.**
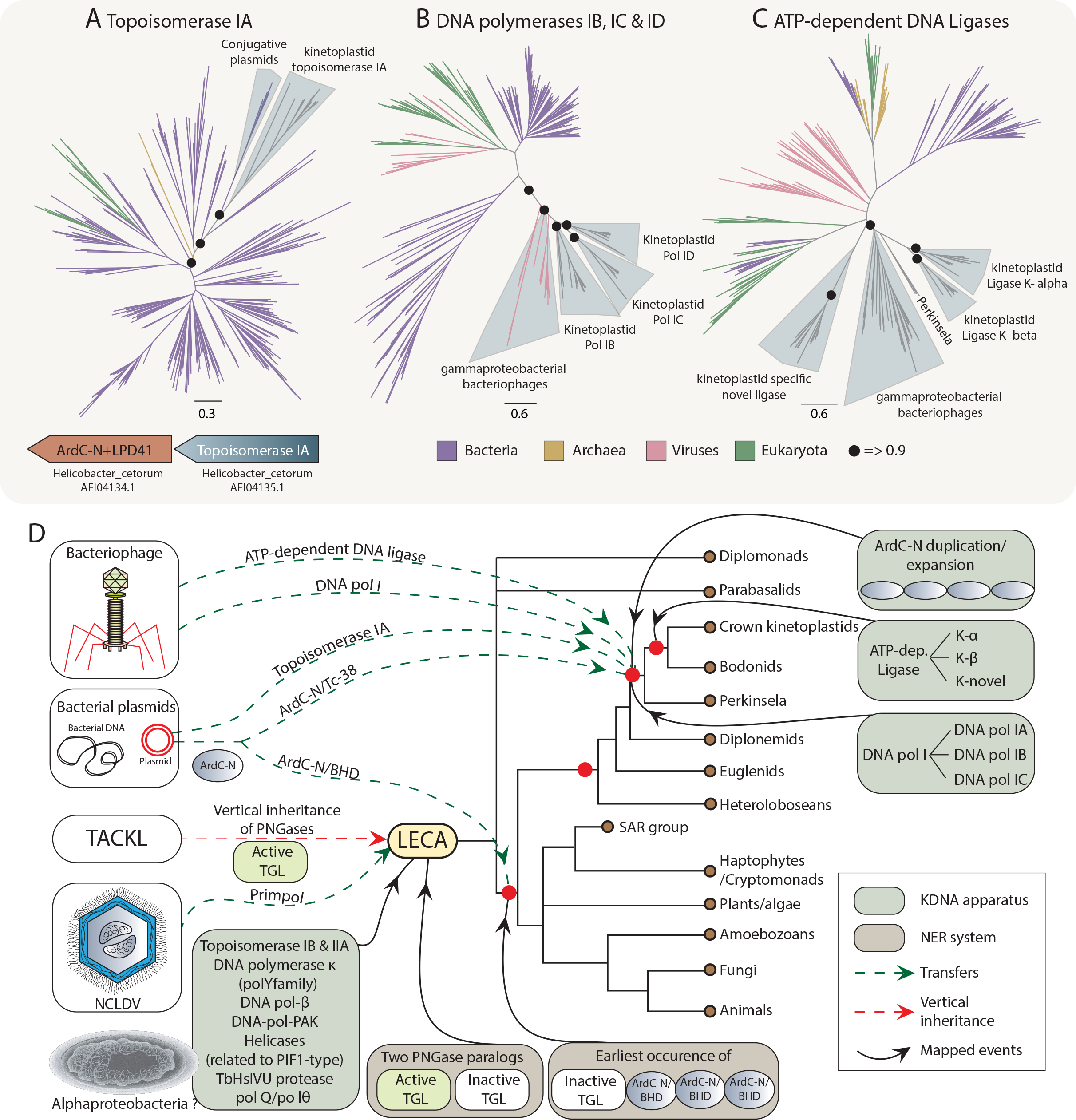
Phylogenetic trees showing key members of the kDNA replication apparatus, namely (A) topoisomerase IA, (B) DNA polymerases IB, IC, and ID, and (C) ATP-dependent DNA ligases, being acquired from plasmids/conjugative elements or bacteriophages. Operonically linked ArdC-N and topoisomerase IA in plasmids/conjugative elements are shown for *Helicobacter cetorum*. D) Schematic diagram showing the summary of the evolutionary events that shaped the kinetoplastid replication and NER systems in eukaryotes. Major evolutionary events are mapped on to a simplified eukaryotic tree. Transfers from plasmids/conjugative elements, bacteriophages and NCLDV sources are highlighted in green dotted arrows. Events related to the kDNA-replication apparatus and the NER-repair system are highlighted in green and brown boxes, respectively and are mapped to the respective nodes on the eukaryotic tree (using curved lines in black and arrowheads).

These are joined by a second category of kDNA replication proteins which either have a deep evolutionary history as mitochondrial replication proteins or function as in eukaryotic nuclear replication. These include two other topoisomerases (DNA topoisomerase IB and IIA) involved in topological manipulations of kDNA [75-77], DNA polymerase κ homologs of the polY family functioning in translesion repair in most eukaryotes [78, 79], two DNA polymerase β (DNA pol-β) enzymes (including DNA pol-β-PAK [80]), six distinct helicases (TbPIF1, TbPIF2, TbPIF4, TbPIF5, TbPIF7, and TbPIF8 [81] related to the eukaryotic dual mitochondrial and nuclear PIF1-type helicase [82], and the TbHsIVU protease [83]. The primary eukaryotic mitochondrial DNA polymerase pol Q/pol θ (Pol IA in *T. brucei*) was retained in kinetoplastids but was apparently relegated to a role in repair [84, 85]. The third category of proteins recruited to kDNA replication includes genuinely kinetoplastid-specific proteins lacking homologs elsewhere. One of these, p93, appears to contribute in a poorly-understood role to minicircle replication [86]. We could trace the provenance of p93 to *Bodo*, however, no homolog was found in *Perkinsela*.

From this survey, it is apparent that the kDNA replication apparatus represents a major reconfiguration of the ancestral mitochondrial replication system inherited by eukaryotes from an alphaproteobacterial ancestor [1, 87, 88]. Based on the mechanistic details of kDNA replication it had been proposed that the precursor of the kDNA circles was perhaps a plasmid harbored within the mitochondrion of the common ancestor of kinetoplastids [22]. The current study allows us to objectively evaluate this proposal and present an evolutionary scenario for the origin of kDNA replication system (Fig 4D). First, our analysis indicates that key components of the kDNA replication system emerged from independent replicons found in bacteria; namely, plasmids/conjugative elements and bacteriophages. There is no evidence for these elements residing in the eukaryotic mitochondrion at a time long after the stem eukaryote, where the endosymbiosis first occurred. However, eukaryotes including kinetoplastids harbor other bacterial endosymbionts (e.g. certain bodonids, strigomonads and novymonads [89-91], which might contain such selfish elements. As the mitochondrial genome degenerated it reached a size comparable to that of these selfish replicons, which code for their own replication proteins. Given that these are likely optimized for the dedicated replication of such small replicons, they in part displaced the ancestral components of the mitochondrial replication system. In parallel, as we had earlier shown [13], central components of the RNA-editing system of the kinetoplast such as the RNA-ligases and end-processing enzymes were also derived from bacteriophage RNA-repair systems. Together with the acquisition of key components DNA replication components, such as the Tc-38 and Topo IA from plasmids, DNA pol I and ATP-dependent ligases from bacteriophages and the primpol from NCLDVs, these probably facilitated the unique structural developments that characterize kDNA. Additionally, they were supplemented by components already present in the ancestral eukaryote as well as lineage-specific innovations.

## CONCLUSIONS

In this study, we show that the N-terminal domain of the anti-restriction protein ArdC of plasmids/conjugative elements was acquired by eukaryotes on two independent occasions and incorporated into two different functional systems. The first acquisition, which happened early in eukaryotic evolution, gave rise to the so-called BHDs of the XPC/Rad4. The second acquisition in the common ancestor of the kinetoplastids gave rise to the Tc-38 like proteins that extensively diversified in the context of the kDNA replication and dynamics. In both instances, the ArdC-N domain underwent multiple duplications within the same polypeptide in eukaryotes to form an extensive DNA-binding interface that could span a substantial length of DNA. In the case of the BHDs, this was accompanied by substantial structural modifications in each of the copies. These findings resolve two outstanding questions regarding very different systems in eukaryotes. One, with more general implications, explains the origin of the eukaryote-specific primary DNA lesion-recognition component in the NER system, several of whose core components were inherited from the archaeal progenitor [43, 44, 92]. We show that a mobile-element-derived DNA-binding protein was re-purposed for this function supplanting the ancestral protein SSB/RPA in this system process. The other acquisition with more specific implications helps explain how the plasmid-like replication and unusual structure of the kDNA of kinetoplastids might have emerged. While in the case of the Tc-38-like proteins their role in kDNA replication is likely to be similar to their ancestral role in mobile-element replication, in the case of XPC/Rad4 there was a shift to a DNA-repair role. In this regard, it might be noted that other DNA-binding domains involved in DNA repair, such as the MutS-I domain, are also shared with the plasmid replication proteins like ArdC-N [8]. In both cases, the incorporation of the ArdC-N domains into eukaryotic systems involved a complex history of physical and/or functional linkage with other components with shared or distinct evolutionary trajectories. In the case of the XPC/Rad4 proteins, they were fused to an inactive TGL related to the active version found in PNGases that catalyze sugar-removal in glycan degradation. Here we establish that these PNGase domains have a deep history in archaea, where they likely performed an ancestral role in the removal of sugar moieties from proteins. The fusion of the TGL domain with the ArdC-N domain was likely a key factor in the emergence of the eukaryote-specific regulation of NER via the ubiquitin-proteasome system. In the case of the kinetoplastid Tc-38-like proteins, we present evidence that the founding member was likely acquired together with a mobile-element-encoded Topo IA which was also incorporated into the kDNA system for topological manipulations of during and after replication. Further, these proteins combined with other acquisitions from bacteriophage and NCLDV sources to constitute the distinctive kDNA replication system.

These findings add to a growing body of case studies that are revealing key aspects of the provenance of eukaryote-specific systems. For instance, the Vrr-Nuc, a nuclease from bacterial selfish-elements, has been recruited into a parallel eukaryotic DNA-repair complex, the Fanconi anemia complex, required for the repair of interstrand cross-links [93, 94]. Strikingly, an associated ssDNA-binding domain, the HIRAN domain, which is also derived from bacterial selfish elements [94-96], has been recruited in eukaryotes for the recognition of the 3’ends of stalled replication forks to facilitate their regression to control DNA damage. In both cases, these domains acquired from the bacterial selfish elements have been physically or functionally linked to components of the ubiquitin-proteasome system to regulate the actual process of DNA repair. Similarly, we had earlier shown that another eukaryotic ssDNA-binding protein involved in recombination, Rad52, had its origin in ssDNA-binding proteins involved in bacteriophage genome recombination [97, 98]. Thus, the sudden emergence of several novel components and whole new systems for the safeguarding of the larger and linearly segmented genomes of eukaryotes appear to have widely recruited pre-adapted catalytic and DNA-binding functions found in prokaryotic selfish elements. This emphasizes the role of selfish elements resident in bacterial endosymbionts, both from the mitochondrial progenitor as well as from others, as major contributors to eukaryote-specific systems. In more general terms, over the years we have presented several lines of evidence for the extensive re-use of domains that ultimately originated in prokaryotic biological conflict systems in eukaryote-specific systems [1, 99]. The opportunities for rapid evolution seen in such conflict systems, which are under intense selective pressures due to their direct effects of fitness, has enabled them to explore a wide structure- and substrate-space thereby furnishing pre-adaptations that could be utilized elsewhere [9].

In conclusion, our findings can help guide specific experiments on DNA-recognition components both in the context of NER and kDNA dynamics. Further, our discovery of bacterial prototypes for some key DNA-recognition components in eukaryotes provides a clear picture of the structural diversity of domains, such as the ArdC-N domain. This could help guide the structural engineering of DNA-binding domains for recognition of specific features such as lesions in DNA.

## Materials and Methods

Iterative sequence profile searches were performed using the PSI-BLAST [100] and JACKHMMER [101] programs against the nr protein database of the National Center for Biotechnology Information (NCBI). Similarity-based clustering implemented in the BLASTCLUST (ftp://ftp.ncbi.nih.gov/blast/documents/blastclust.html) was used for classification/paralogous grouping and culling of nearly identical sequences. The length (L) and score (S) threshold parameters were variably set between 1.2 to .3 to obtain clusters at different thresholds. Profile-Profile searches were run using the HHpred program either against the PDB or Pfam databases [28, 29]. Multiple sequence alignments were built using the KALIGN [102] and GISMO [103] programs, followed by manual adjustments based on profile-profile and structural alignments. Secondary structures were predicted with the JPred [104] program. Phylogenetic relationships were derived using an approximate maximum likelihood (ML) method as implemented in the FastTree program [105]: corresponding local support values were also estimated as implemented. To increase the accuracy of topology in FastTree, we increased the number of rounds of minimum-evolution subtree-prune-regraft (SPR) moves to 4 (-spr 4) as well as utilized the options -mlacc and -slownni to make the ML nearest neighbor interchanges (NNIs) more exhaustive. Phylogenetic tree topologies were also derived using ML methods based on the edge-linked partition model as implemented in the IQ-TREE software [106]: branch supports were obtained using the ultrafast bootstrap method (1000 replicates). Gene neighborhoods was retrieved by a Perl script that extracts the upstream and downstream genes of the query gene and uses BLASTCLUST to cluster the proteins to identify conserved gene neighborhoods. Position-wise Shannon entropy (H) was computed using a custom script written in the R language using the equation

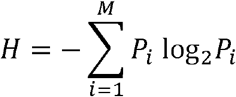

where M is the number of amino acid types and P is the fraction of residues of amino acid type *i*. The Shannon entropy for any given position in the MSA ranges from 0 (absolutely conserved one amino acid at that position) to 4.32 (all 20 amino acid residues equally represented at that position). Structural visualization and manipulations were performed with the PyMol (http://www.pymol.org) program. The in-house TASS package, which comprises a collection of Perl scripts, was used to automate aspects of large-scale analysis of sequences, structures, and genome context.

## Declarations

### Ethics approval and consent to participate

Not applicable.

### Consent for publication

Not applicable.

### Availability of materials

All data analyzed in this paper were downloaded from searches against publicly available databases. All identified sequences and complete data associated with the manuscript is also available at the FTP site: ftp://ftp.ncbi.nih.gov/pub/aravind/ardcn/ardcn.html.

## Authors’ contributions

L.A, A.M.B and L.M.I conceived the project and directed its management. A.K, A.M.B and L.A performed all computational analyses and analysed the data. A.K and A.M.B wrote the first draft of the manuscript. A.K prepared all the figures and tables and supplementary material. L.A edited the manuscript to prepare the final version. All authors read and approved the final manuscript.

## Acknowledgements

This research was supported by Intramural Research Program of the National Institutes of Health (NIH), National Library of Medicine (NLM).

